# Enhanced Thompson Sampling by Roulette Wheel Selection for Screening Ultra-Large Combinatorial Libraries

**DOI:** 10.1101/2024.05.16.594622

**Authors:** Hongtao Zhao, Eva Nittinger, Christian Tyrchan

## Abstract

Chemical space exploration has gained significant interest with the increase in available building blocks, which enables the creation of ultra-large virtual libraries containing billions or even trillions of compounds. However, the challenge of selecting most suitable compounds for synthesis arises, and one such challenge is hit expansion. Recently, Thompson sampling, a probabilistic search approach, has been proposed by Walters *et al*. to achieve efficiency gains by operating in the reagent space rather than the product space. Here, we aim to address some of its shortcomings and propose optimizations. We introduce a warmup routine to ensure that initial probabilities are set for all reagents with a minimum number of molecules evaluated. Additionally, a roulette wheel selection is proposed with adapted stop criteria to improve sampling efficiency, and belief distributions of reagents are only updated when they appear in new molecules. We demonstrate that a 100% recovery rate can be achieved by sampling 0.1% of the fully enumerated library, showcasing the effectiveness of our proposed optimizations.

---

High attrition rates plague drug discovery at all stages and can often be traced back to the quality of the chemical leads, driving the need to explore a greater chemical space.^1^ Parallel to high-throughput screening,^2^ DNA-encoded library technology^3^ and fragment-based lead discovery,^1^ virtual screening offers a cost-effective and time-efficient means for hit identification, typically applied to collections of a few millions molecules.^4^ As the library grows, better-fitting molecules are found and the score improves log-linearly with the library size.^5, 6^

The ultra-large virtual libraries, on the scale of billions or even trillions, can be constructed from much smaller sets of reagents. The number of available building blocks has grown to hundreds of thousands. For instance, the three-component Niementowski quinazoline reaction yields a fully enumerated library of 94 million products, for a reagent pool of having 376 aminobenzoic acids, 500 primary amines, and 500 carboxylic acids (**Figure 1**).^7^ To screen ultra-large virtual libraries in a brute-force fashion is costly and requires special computing infrastructures as well.^8, 9^ Harnessing the combinatorial nature of ultra-large libraries, the V-SYNTHES approach decomposes the library into sets of scaffolds and synthons from which a minimal enumeration library is constructed by attaching a single synthon to a scaffold at one R-group position.^10, 11^ The remaining attachment points on the scaffold are capped with methyl or phenyl groups. The most promising fragments are identified upon docking, and their capped R-groups are iteratively enumerated until the molecules are fully elaborated.

**Figure 1.**
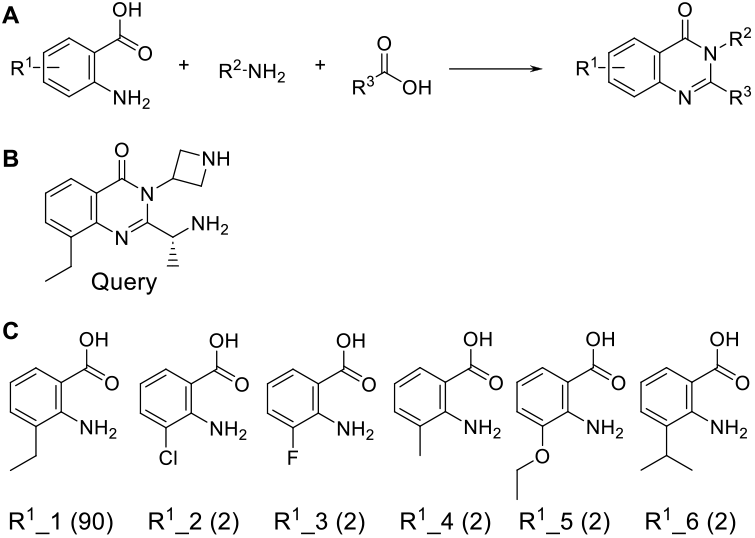
The combinatorial library based on Niementowski quinazoline synthesis. (**A**) The three-component reaction scheme; (**B**) the query molecule and (**C**) the six R^1^ reagents in the 100 top-scoring hits from an exhaustive similarity search of the 94 million molecules enumerated with 376 R^1^, 500 R^2^ and 500 R^3^ reagents. The number in the bracket is the frequency of occurrence in the top 100 molecules.

Alternatively, Thompson sampling, a probabilistic search approach, has recently been proposed to achieve efficiency gains by working in the reagent space rather than the product space.^7^ It can be applied to hit expansion in drug discovery. This exploitation is focused on a specific chemical space and measures the relevance of reagents by a scoring or similarity function. Remarkably, upon screening 0.1% of the library, it discovers 90 out of the 100 top-scoring molecules, which have been identified from an exhaustive search of the above 94 million quinazoline molecules by measuring their Tanimoto similarity to a query molecule. However, the recovery rate has plateaued at 90% even by concatenating the results from multiple searches. Notably, it identifies only the most prevalent one of the six R^1^ building blocks contained in the 100 top-scoring molecules (**Figure 1**). In this letter, we show that the stochastic roulette wheel selection can improve the Thompson sampling efficiency and lead to a 100% recovery rate upon screening 0.1% library, in comparison with the original greedy selection approach.

Thompson sampling lies in Bayesian inference by approximating the actual likelihoods as normal distributions which are justified by the central limit theorem.^12^ Consider a single observation *y*_*i*_ of scores associated with a reagent *i* from a normal distribution parameterized by an unknown mean *θ*_*i*_ and known variance *σ*^2^, the sampling distribution is

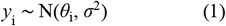

Take *θ*_*i*_ as an exponential of a quadratic form, the family of conjugate prior densities is

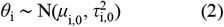

where *μ*_i,0_ and *τ*_i,0_ are the prior mean and variance for reagent *i*, respectively. Given a sample of independent and identically distributed observations *y* = (*y*_1_, …, *y*_n_) for reagent *i*, its posterior mean and variance are

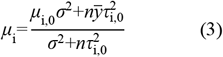

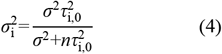

Where 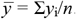. The posterior predictive distribution 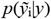 of a future observation 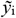 for reagent *i* is

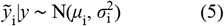

The initial belief distributions are produced from a warmup procedure.^7^ In a warmup cycle, the reagents of each reaction component are placed into a corresponding column of the matrix ***M***_m×n_ by repeating themself until that column is filled up, wherein *m* is the number of components in the reaction and *n* is the largest number of reagents among all components. This ensures that each reagent will be selected at least once in the minimum *n* molecules. Each row of ***M***_m×n_ is then selected for making and scoring the molecules. The warmup cycle is repeated three times, with the reagents being shuffled each round. The empirical mean 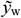 and variance *σ*_w_ are calculated from all scores observed during warmup. The known variance *σ*^2^ as well as the prior mean *μ*_i,0_ and variance *τ*_i,0_^2^ for all reagents are set as follows^7^

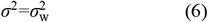

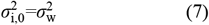

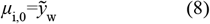

Following the warmup, the posterior distribution of each reagent is updated with its scores seen during warmup by **Eq. 3** and **4**. Next, repeat the search cycle which consists of the following steps:

**Step 1**: Sample one 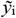 for each reagent *i* by **Eq. 5**, and convert it into a probability *p*_i_ among the reagents of its corresponding component

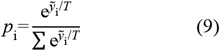

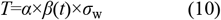

where the summation runs over all reagents in that component; *T* is the Boltzmann temperature parameter; *α* is a scaling factor which is positive if higher *y* is preferred and negative otherwise; *β*(*t*) is a time-dependent dumping coefficient and *t* is the number of search cycles performed. Throughout the manuscript, *α* = *β*(*t*) = 1.

**Step 2**: Sample *n* reagents from each reaction component by roulette wheel selection according to their probability distribution. Similar to the warmup procedure, *n* molecules are selected for making. If the molecules are new, make and score them; otherwise, skip the subsequent step for those made in the previous rounds.

**Step 3**: Update the posterior distributions of the selected reagents. The posterior mean of reagent *i* for a three-component reaction converges according to

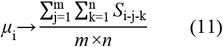

wherein *m* and *n* are the number of reagents in the other two components, respectively; and *S*_i-j-k_ is the score of the molecule made from reagents *i, j* and *k*. Its variance converges to *σ* _w_ ^2^/(*m*×*n*+1).

The search is aborted if one of the two stop criteria has been satisfied: the maximum number of iterations being reached or failure to sample a new molecule for the given number of search cycles in a row (default 100).

To illustrate its sampling efficiency, the new method is applied to identify molecules in the abovementioned quinazoline combinatorial library most similar to the query (**Figure 1**). The warmup scores of the 1500 molecules resulting from 3 cycles show a normal distribution of the Tanimoto coefficients with a fitted mean of 0.2 and standard deviation of 0.04 (**Figure 2**). Following the warmup procedure, the posterior distributions of the six R^1^ reagents have a mean of 0.326 and standard deviation of 0.02 for R^1^_1, 0.291 ± 0.016 for R^1^_2, 0.229 ± 0.018 for R^1^_3, 0.249 ± 0.018 for R^1^_4, 0.247 ± 0.02 for R^1^_5, 0.284 ± 0.016 for R^1^_6. The probability of selecting R^1^_2 or R^1^_3 over R^1^_1 by the greedy selection (i.e., taking the maximum of the sampled values)^7^ is 8% and 0.01%, respectively, calculated by

**Figure 2.**
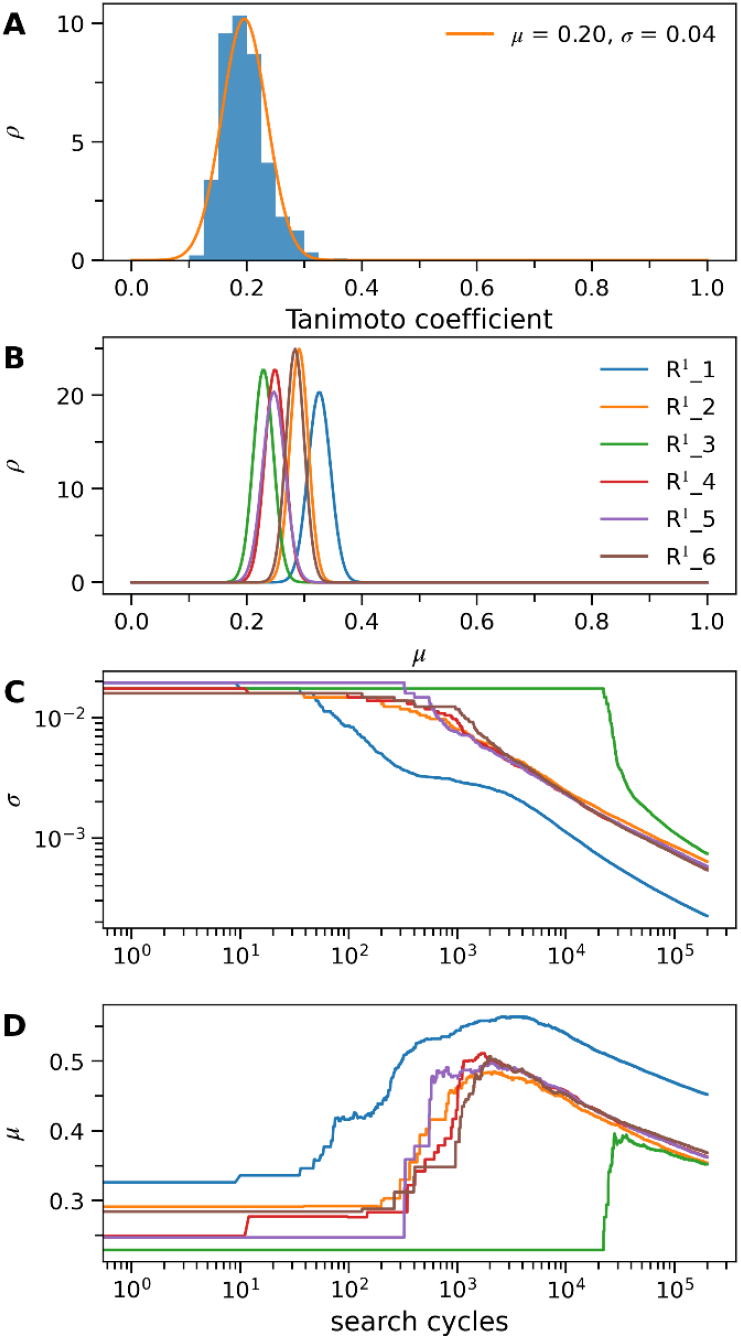
Prior and posterior distributions of the six R^1^ reagents. (**A**) Distribution of the warmup scores (i.e., Tanimoto coefficients to the query using the 2048-bit ECFP4 fingerprint); (**B**) posterior distributions of the six R^1^ reagents following the warmup; and evolution of their standard deviations (**C**) and mean values (**D**) over the search with one molecule per cycle. The color scheme is consistent from **B** to **D**.

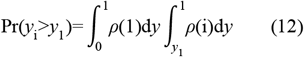

where *i* indicates one of the other five R^1^ reagents. Note that R^1^_2 is closest to R^1^_1 while R^1^_3 is furthest with regard to the initial posterior distributions of the six R^1^ reagents (**Figure 2B**). Consequently, R^1^_1 would be prioritized and matched with high-score reagents from the other two components, shifting its mean toward the high-score region (**Figure 2D**). Concomitantly, its posterior variance decreases quickly as it is inversely proportional to the number of times being sampled (**Figure 2C**). Combined, it explains the failure to sample the other five R^1^ reagents by the greedy selection.^7^

In comparison, the probability by roulette wheel selection is 47.4%, 19.4% and 4.0% for selecting R^1^_1, R^1^_2 and R^1^_3 among the six R^1^ reagents, respectively. Note that the temperature parameter *T* is controllable by **Eq. 10**. As shown in **Figure 2C**, R^1^_1 has been selected in the first 10 search cycles, followed by R^1^_2, R^1^_4, R^1^_5 and R^1^_6 within a few hundreds of cycles; while R^1^_3 is selected for the first time at around 20 000 cycles. Note that if a reagent is not selected during a search cycle, its variance remains unchanged. Intriguingly, the posterior mean of all six R^1^ reagents increase rapidly to a peak upon being sampled, and then decrease gradually, which is due to an unchanged belief distribution for reagents of a molecule that has been sampled previously. After exhausting highscore reagents from the other two components, the pairing of a high-score reagent with low-score ones over the search drives its mean toward convergence in the low-score region. This would then increase sampling of initially low-score but intrinsically high-score reagents such as R^1^_3. For example, the difference in the mean among the five R^1^ reagents (2-6) after around 100, 000 cycles is much smaller than that at the beginning of the search, consistent with their equal frequency of occurrence in the 100 top-scoring hits (**Figure 1**).

The performance of recovering the 100 top-scoring hits averaged over the 10 replicates is shown in **Figure 3A**. Around 60% of the 100 top-scoring hits could be retrieved by screening only 0.01% of the library, and all the 100 hits can be discovered upon screening 0.1% of the library. The efficiency of sampling new molecules, measured by the ratio of the unique molecules to the total, decreases sharply to around 40% within 0.01% of the library being screened, suggesting that initially high-score reagents are oversampled (**Figure 3B**). Afterwards, the sampling efficiency increases quickly when initially low-score but intrinsically high-score reagents begin to be selected. The sampling efficiency is around 80% at the 0.1% of the library when all the 100 hits have been recovered. It continues to increase as the mean of those high-score reagents start to drop, suggesting that it navigates into the largely uncharted chemical space rather than being trapped in the already mapped-out regions. There is no significant difference in the performance between sampling one and three molecules per cycle. However, it is computationally efficient to sample three molecules since 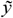 of all reagents only need to be sampled once. As *in silico* make and the subsequent similarity comparison is relatively fast, the step of sampling 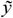 is rate-limiting. The computational time by sampling three molecules is roughly one-third of that by sampling one molecule per search cycle. In case of time-consuming scoring methods such as docking, sampling multiple molecules per search cycle allows for parallelization which is critical to dock even 0.1% of ultralarge libraries.

**Figure 3.**
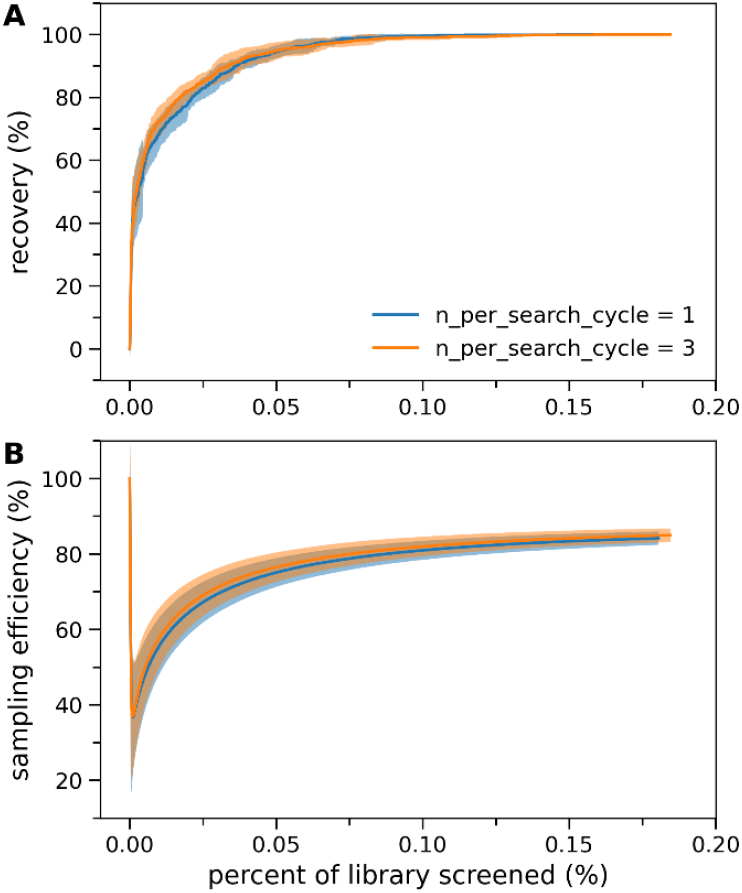
Performance of the enhanced TS sampling averaged over the 10 replicates. (**A**) Recovery of the 100 top-scoring hits and (**B**) efficiency of sampling new molecules.

In summary, the Thompson sampling efficiency can be enhanced by roulette wheel selection and the passive penalization of high-score reagents over the search. Furthermore, the roulette wheel selection allows for sampling multiple molecules per search cycle, which is critical to parallelization of scoring multiple molecules. In addition, we introduce a new warmup approach to ensure that each reagent will be selected within a minimum number of molecules.

## ASSOCIATED CONTENT

### Data Availability Statement

The source code of implementing the enhanced TS is available at https://github.com/WIMNZhao/TS_Enhanced.

## AUTHOR INFORMATION

### Notes

H.Z., E.N. and C.T. are employees of AstraZeneca, and may own stock or stock options in AstraZeneca.

## ACKNOWLEDGMENT

The authors are grateful to Dr. Patrick Walters for sharing their TS codes and the quinazoline library publicly.

---

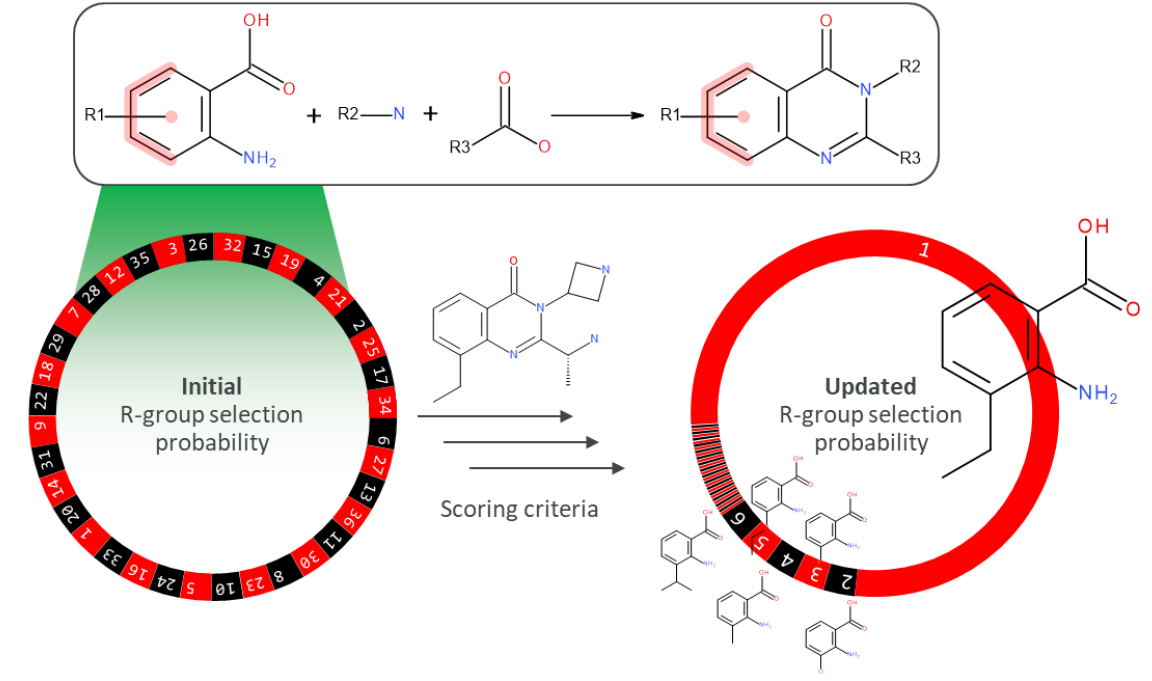

